# Binding mechanism underlying FIH-1 suppression caused by the N-terminal disordered region of Mint3

**DOI:** 10.1101/2021.05.10.443412

**Authors:** Tensho Ten, Satoru Nagatoishi, Masaru Hoshino, Yoshiki Nakayama, Motoharu Seiki, Takeharu Sakamoto, Kouhei Tsumoto

## Abstract

Mint3 is known to enhance aerobic ATP production, known as the Warburg effect, by binding to FIH-1. Since this effect is considered to be beneficial for cancer cells, the interaction is a promising target for cancer therapy. However, previous research has suggested that the interacting region of Mint3 with FIH-1 is intrinsically disordered, which makes investigation of this interaction challenging. Therefore, we adopted a physicochemical approach that combined thermodynamic studies with structural analyses in solution, to clarify the binding mechanism. First, using a combination of CD, NMR, and hydrogen/deuterium exchange mass spectrometry (HDX-MS), we confirmed that the N-terminal half, which is the interacting part of Mint3, is mostly disordered. Next, using isothermal titration calorimetry (ITC), we revealed that the interaction of Mint3 and FIH-1 produces an enormous change in enthalpy and entropy. The profile is consistent with the model that the flexibility of disordered Mint3 is drastically reduced upon binding to FIH-1. Moreover, we performed a series of ITC experiments with several types of truncated Mint3s, an effective approach since the interacting part of Mint3 is disordered, and identified 78-88 as a novel core site for binding to FIH-1. The truncation study of Mint3 also revealed the thermodynamic contribution of each part to the interaction, where one contributes to the affinity (Δ*G*), while the other only affects enthalpy (Δ*H*), by forming non-covalent bonds. This insight can serve as a foothold for further investigation of IDRs and drug development for cancer therapy.

## Introduction

Cancer is a major cause of death worldwide. Thus, the development of its treatment is of great interest in various research areas, including molecular targeted therapies. Although the types of cancer vary widely and a large part of the complex mechanism remains unclear, some common factors or phenomena could serve as promising targets for drug development. “Warburg effect” is one of the phenomena that are considered to play a pivotal role in the survival and proliferation of tumor cells. It is characterized by increased aerobic glycolysis, even in the presence of abundant oxygen, which is frequently observed in malignant tumor cells. Since the glycolytic approach of energy production is less efficient in terms of ATP production than oxidative phosphorylation, which is the primary pathway of ATP production in most normal cells, the preference for aerobic glycolysis is assumed to have some implications on the differentiation between cancer cells and normal cells. Although the benefits of Warburg effect for cancer cells are still unclear, numerous proposals have been made in the past decades, based on intensive research (1, 2). The possible explanations are an adaptation to hypoxia, which is the condition that most cancer cells confront, and/or acceleration of production of building blocks required for cell proliferation, which are provided as byproducts of glycolysis. The mechanisms of Warburg effect are also being gradually revealed along with the investigation of these functions. Hypoxia-inducible factor-1 (HIF-1) is a key transcription factor that regulates the expression of genes necessary for cellular responses to hypoxic conditions. The major examples of the proteins expressed from these genes include vascular endothelial growth factor, erythropoietin, glucose transporters, and several glycolytic enzymes such as phosphoglycerate kinase 1.

HIF-1 is composed of α and β subunits, and its transcriptional activity is regulated by factor inhibiting HIF-1 (FIH-1), an asparaginyl hydroxylase that prevents HIF-1α binding to p300/CBP through modification of the Asn803 residue. Since the enzymatic activity of FIH-1 requires Fe (II) and α-ketoglutaric acid (αKG), the regulation of HIF-1 is known to be oxygen-dependent. However, Sakamoto and Seiki identified another factor, Mint3, a member of the X11 protein family, which controls the suppression activity of HIF-1 by FIH-1 (3). Previous research by Sakamoto and Seiki showed that Mint3 binds to FIH-1 through its N-terminal portion and inhibits the regulatory activity of FIH-1 by competing with HIF-1α (3). Accordingly, the protein-protein interaction (PPI) between Mint3 and FIH-1 is one of the critical pathways that enhances the activity of HIF-1, which is followed by a metabolic effect that is beneficial for cancer cells. Therefore, Mint3-FIH-1 PPI is a promising target for the development of cancer therapies.

Additionally, a previous research has implied that the N-terminal region of Mint3, which is the binding site of FIH-1, is intrinsically disordered (3). Although it is traditionally assumed that the function of a protein originates in its three-dimensional structure, many intrinsically disordered proteins (IDPs) and intrinsically disordered regions (IDRs) have recently been identified to be biologically functional (4–7). Moreover, most of these biological functions are activated by PPIs between IDPs/IDRs and their binding partners. Therefore, the hypothesis that the intrinsically disordered part of Mint3 inhibits FIH-1 is feasible and attractive enough to be validated. However, owing to the difficulties in analyzing the structural dynamics of IDPs/IDRs, clarification of the molecular mechanisms of their PPIs is still challenging. Based on its pharmaceutical importance as a promising target of cancer therapy and the scientific interest in the PPIs of IDPs/IDRs, we performed a series of physicochemical analyses to describe the binding mechanism of Mint3 to FIH-1 at a molecular level.

## Results

### Intrinsically disordered N-terminal region of Mint3

Mint3, also known as X11γ/APBA3, belongs to the X11 protein family. Although the remaining two members, Mint1/X11α/APBA1 and Mint2/X11β/APBA2, are neuronal proteins, Mint3 is ubiquitously expressed (8). All three X11s have a conserved C-terminal half that is composed of a phosphotyrosine-binding (PTB) domain, which is known to be the binding site of amyloid precursor protein (APP), and two PDZ domains (Figure 1a). However, their N-terminal halves are variable. Although both Mint1 and Mint2 have an N-terminal binding site to munc18-1, which is a neural protein involved in synaptic vesicle exocytosis, Mint3 does not have one (3, 9). Instead, the N-terminal region (1-214) of Mint3 is reported to mediate binding to FIH-1 (Figure 1a), although it is supposed to be intrinsically disordered according to its primary structure (3). To characterize the structure of the involved region, we expressed the N-terminal fragment (1-214) of Mint3 (hereafter called Mint3NT) and performed a series of structural analyses in solution.

**Figure 1:**
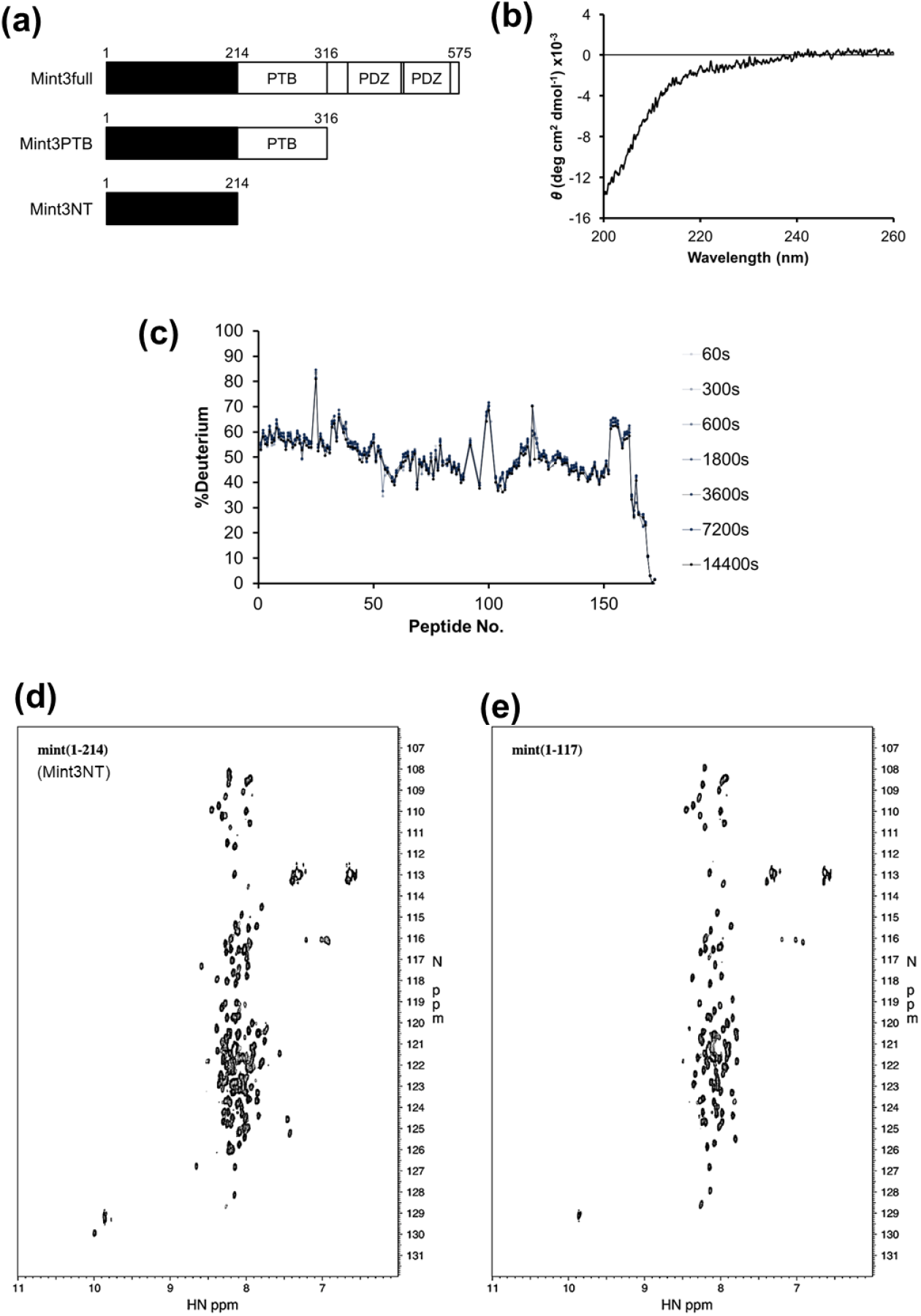
Molecular information of Mint3. **(A)** Mint3 constructs used in this study. **(B)** CD spectrum of Mint3NT. **(C)** HDX-MS chart of Mint3NT. **(D)** NMR spectra of Mint3NT and Mint3 (1-117).

Circular dichroism (CD) experiments were performed to measure the composition of the secondary structure of Mint3NT. However, significant peaks that originate in secondary structures, especially α-helices and β-sheets, could not be observed (Figure 1b), thus suggesting that the isolated N-terminus of Mint3 is mostly composed of a random coil (10). Following that, differential scanning calorimetry (DSC) was performed to analyze the protein folding of Mint3NT. No significant peak caused by the collapse of the higher-order structure was observed in the thermal denaturation experiment (Figure S1).

To determine the structural molecular details of the N-terminal regions of Mint3, hydrogen/deuterium exchange mass spectrometry (HDX-MS) and nuclear magnetic resonance (NMR) spectroscopy were conducted. In the HDX-MS analysis, the exchange events are mediated by the conformational fluctuation of the protein; thus, the exchange rate of amide hydrogen is mostly influenced by the formation of a hydrogen bond (11, 12). Therefore, the amide hydrogen of an extended/flexible region will be replaced by solvent deuterium at a higher rate, as compared to that of a folded/rigid region. In fact, the exchange rates of whole Mint3NT were observed as an immediate saturation in the deuteration level, suggesting an intrinsically disordered state in solution (Figure 1c). NMR analysis was carried out for Mint3NT and its truncated mutant (1-117). Mint3NT provided a spectrum with poorly dispersed resonance peaks, particularly along the ^1^H-axis, in which most of the peaks appeared in the disordered region (8.5–7.5 ppm). However, the linewidth of each peak was very sharp, and the resonances derived from the sidechains of the three tryptophans could be observed around (10 ppm, 130 ppm) separately (Figure 1d). The truncated mutant (1-117) also exhibited a similar spectrum, and the peak of a single tryptophan was observed in the same region as seen in case of Mint3NT (Figure 1e). All these results suggested that Mint3NT is mostly unstructured in solution. With several exceptional residues between 110-117, the resonance peaks in the two spectra mostly overlapped. Thus, it was clear that isolated Mint3NT is mostly disordered in solution. Also, we considered that truncation of Mint3NT is an effective methodology for the following detailed interaction analysis.

### Physicochemical properties of the interaction

Sakamoto *et al*. identified Mint3NT as a construct that is sufficient for binding to FIH-1 in immunological assays (3). We performed a couple of thermodynamic analyses on the interaction between Mint3 and FIH-1 using isothermal titration calorimetry (ITC), to confirm the interaction quantitatively. The Mint3 constructs used in these experiments were Mint3PTB (1-361), which contains the PTB domain within the C-terminal half, and Mint3NT. Exothermic interactions were observed for both the Mint3s. Moreover, the thermodynamic parameters [the binding free energy (Δ*G*), binding enthalpy (Δ*H*), and binding entropy (–*T*Δ*S*)] were the same for Mint3PTB and Mint3NT, within the error range (Figures 1a, 2a-b). The PTB domain did not contribute significantly to Mint3NT binding. Thus, we regarded that the Mint3NT construct was of sufficient length in the context of binding to FIH-1. Of note, the binding of Mint3NT to FIH-1 produced an enormous change in enthalpy (Δ*H* = –32.2 ± 0.7 kcal.mol^-1^) and entropy (– *T*Δ*S* = –23.6 ± 0.8 kcal.mol^-1^) (Figure 2b). This phenomenon can be interpreted as follows: (i) the large negative Δ*H* suggests the formation of a large number of intra- and inter-molecular bonds. (ii) The large positive –*T*Δ*S* reflects the loss of conformational flexibility of Mint3NT through the interaction. Additionally, the binding stoichiometry (*N* value) for Mint3NT was estimated as 0.48 ± 0.02. Since in the ITC experiment, Mint3 and FIH-1 were loaded into the syringe and cell, respectively, this number suggests that the binding molar ratio between Mint3 and FIH-1 is one to two. To confirm the oligomerization state of FIH-1, size-exclusion chromatography (SEC) and SEC-multi-angle laser scattering (SEC-MALS) measurements were conducted. In the SEC measurement, the chromatogram showed a single peak at the elution volume of the molar mass of the FIH-1 dimer. The SEC-MALS measurement also provided a chromatogram with a single peak, and the measured molar mass was approximately 80 kDa, which is consistent with that of the FIH-1 dimer (Figure S2). Notably, although the concentration of FIH-1 used in SEC-MALS (∼70 µM) was approximately 10-times greater than that of SEC (∼7 µM), the results suggested that most FIH-1 forms a dimer under both the conditions. Considering that the concentration of FIH-1 used in the ITC measurement (30 µM) was between these two concentrations, it was clear that FIH-1 presented as a dimer uniformly. Therefore, the stoichiometry can be interpreted to indicate that one Mint3 binds to a dimer of FIH-1.

**Figure 2:**
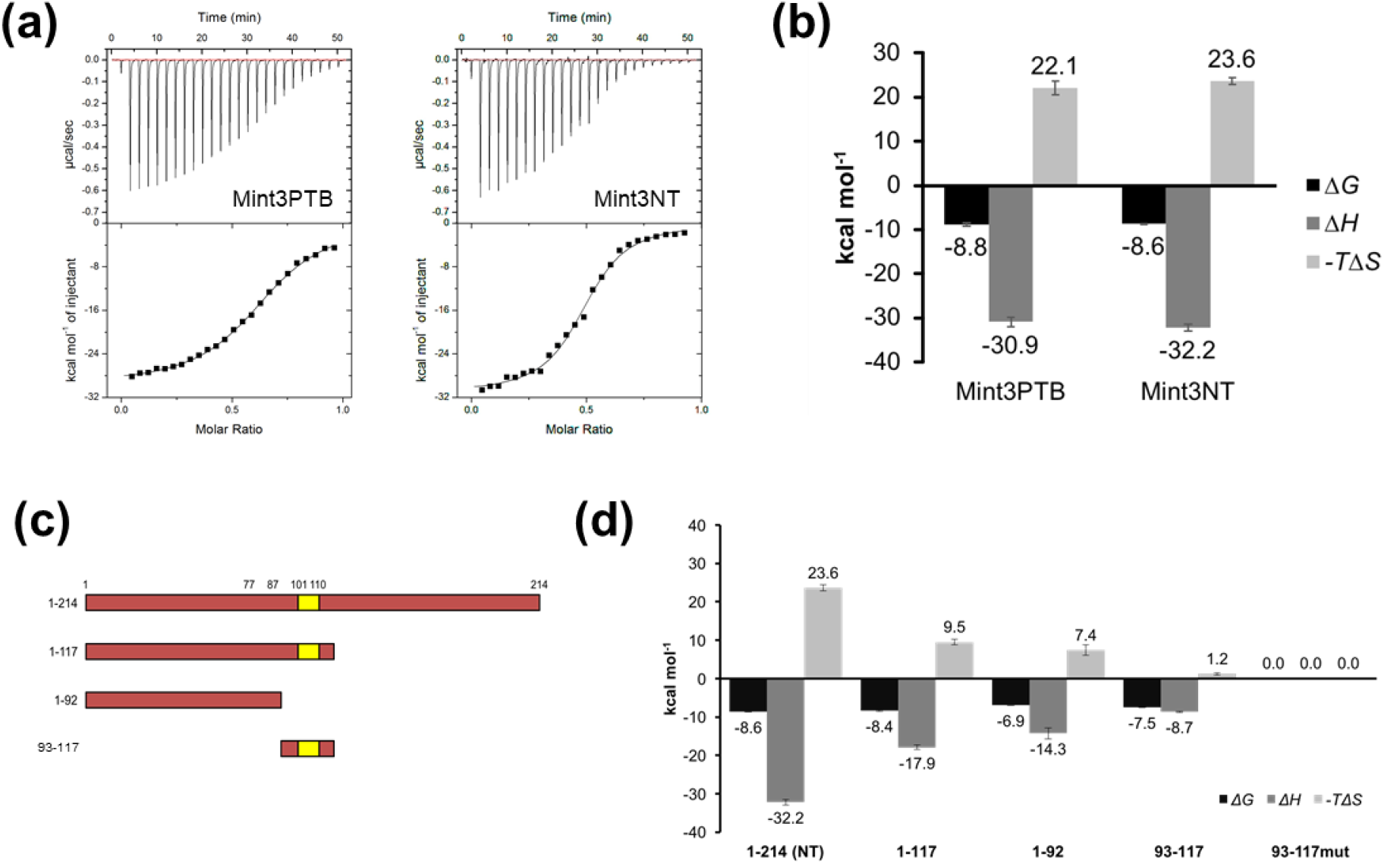
**(A)** ITC profiles of Mint3PTB and Mint3NT binding to FIH-1. **(B)** Thermodynamic parameters of the interactions between FIH-1 and Mint3. **(C)** Deletion mutants of Mint3NT. **(D)** Thermodynamic parameters of the interactions between FIH-1 and deletion mutants of Mint3NT.

The above mentioned structural analysis depicted the intrinsically disordered state of isolated Mint3NT, and the thermodynamic parameters of the interaction suggested a huge loss of conformational flexibility for Mint3NT in complex with FIH-1. CD analysis was further performed to determine whether Mint3NT forms secondary structures through this interaction. We compared the spectrum obtained from the Mint3NT-FIH-1 complex with a theoretical spectrum generated by merging the spectra of isolated Mint3NT and FIH-1 (Figure S3). As both the spectra were similar, Mint3NT did not seem to form secondary structures through binding to FIH-1, suggesting that a significant conformational change was not observed, at least at the level of the secondary structures.

### Thermodynamic/kinetic contribution of each part of Mint3NT to the interaction

In a previous study, the 101-110 region within Mint3NT was identified as a core region responsible for its binding to FIH-1 (13). Accordingly, it is possible that Mint3NT can be divided into the binding and non-binding sites. To clarify the contribution of each part of Mint3NT, especially in quantitative terms, we performed a series of thermodynamic analyses for several truncated and mutated versions of Mint3NT using ITC (Figure 2c-d). Some constructs (1-117, 1-92, 93-117, Mint3NTmut) were observed in the exothermic reactions. The *N* values of these truncated mutants were also around 0.5, suggesting that they bind to a dimer of FIH-1, in a similar manner to Mint3NT.

A short peptide of Mint3 (93-117) containing the predicted core region (101-110) was observed in the ITC measurement, and its Δ*G* was calculated to be –8.6 ± 0.1 kcal mol^−1^, based on the measured equilibrium dissociation constant (*K*_D_ = 3.0 ± 0.6 µM). Next, when alanine mutations were introduced into the peptide 93-117 (<u>GLL</u>SA<u>E</u>AGRD → <u>AAA</u>SA<u>A</u>AGRD, 93-117mut), the interaction was found to be completely disrupted. Notably, the introduction of the same mutations into whole Mint3NT (Mint3NTmut) weakened the binding (*K*_D_ = 1.31 µM, Δ*H* = –13.9 kcal.mol^-1^, –*T*Δ*S* = 5.9 kcal.mol^-1^), but did not disrupt the interaction completely (Figure S4). Thus, we considered that the predicted region (101-110) was just one of the binding sites.

After confirming that Mint3 has a binding site at 101-110, further truncation analyses were conducted. First, the N-terminal half (1-117) of Mint3NT, which contains the binding site (101-110), also exhibited significant exothermic interaction with FIH-1. The measured *K*_D_ was 0.65 ± 0.07 µM and the calculated Δ*G* was the same (within error) with Mint3NT. Notably, the measured Δ*H* for the N-terminal half of Mint3NT was approximately half of that for Mint3NT (Figure 2c-d). Next, a Mint3 fragment (1-92) that lacked the binding site (101-110) was constructed. Remarkably, although the fragment (1-92) displayed a significant decrease in exothermic interaction, it retained its binding ability to FIH-1 (Figure 2c-d). Therefore, we hypothesized that the N-terminal half (1-117) of Mint3NT is sufficient to generate affinity (Δ*G*), and that the C-terminal half (111-214) does not contribute to Δ*G*, but contributes to Δ*H*. Furthermore, there is at least one more binding site in the 1-92 region of Mint3. Thus, we found some binding mechanisms in the ITC experiments, as follows: i) One dimer of FIH-1 binds to one Mint3NT monomer and ii) there are other interaction sites in Mint3NT, apart from Mint3 (93-117).

### Role of the dimer interface of FIH-1

According to the binding stoichiometry measured with ITC, Mint3NT binds to a dimer of FIH-1. Thus, we hypothesized that the pivotal binding site of FIH-1 is located around the dimer interface, from the standpoint of symmetry. Although FIH-1 monomers strongly interact with each other by meshing their hydrophobic C-terminal helices, the substitution of Leu340 for arginine is known to disrupt the dimerization (14). Hence, a single point mutant FIH-1L340R was constructed to examine the role of the dimer interface (Figure S5). As expected, FIH-1L340R was mostly eluted in the monomeric fraction in SEC (Figure 3a).

**Figure 3:**
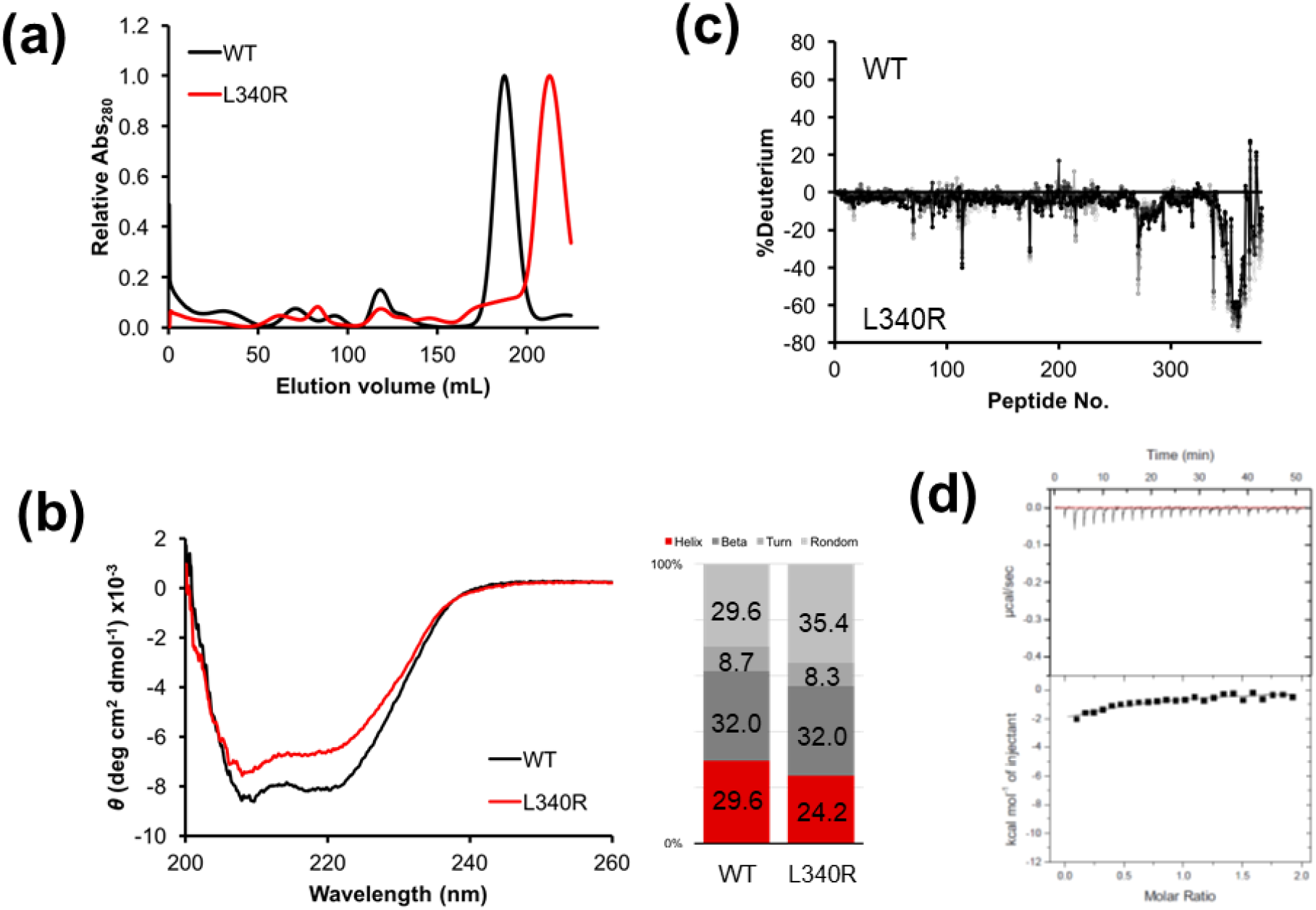
Molecular and interaction data of Mint3NT L340R. **(A)** SEC profiles of Mint3NT and L340R. **(B)** CD spectra of Mint3NT and L340R. **(C)** Differences between HDX-MS charts for Mint3NT and L340R. **(D)** ITC profile of the interaction between Mint3NT L340R and FIH-1.

To examine the conformational identity between the wild-type and monomeric mutant, CD measurements and differential scanning fluorometry (DSF) were conducted. In CD measurements, a recession of α-helicity was observed for the monomeric mutant. In quantitative terms, α-helix decreased by 5.4% and random-coil increased by 5.8%, whereas β-sheet and turn were practically unchanged. These results suggest that the mutation may disturb the structure of FIH-1, especially the α-helices (Figure 3b). In DSF, the monomeric mutant showed lower thermal stability than the wild-type, and notably, the baseline level of the monomeric mutant was higher than that of the wild-type (Figure S6). Because SYPRO^®^ Orange, which is a protein-dye used in the DSF experiment, binds non-specifically to the hydrophobic surface and emits fluorescence, the elevated baseline is likely to represent the exposure of the hydrophobic dimer interface of FIH-1L340R. Additionally, HDX-MS was applied to depict the structural differences between the wild-type and monomeric mutant in detail. The difference in the deuteration level was observed only around the C-terminal α-helices, namely the dimer interface (Figure 3c). According to the results of CD, DSF, and HDX-MS, the substitution of Leu340 for arginine can interrupt the dimerization, by loosening the interfacial α-helices, and the effect on the structure is limited to the periphery of the dimer interface.

Subsequently, binding analysis was performed using ITC, and significant disruption of the binding to Mint3NT was observed for FIH-1L340R. Notably, the binding free energy (Δ*G*) decreased significantly (Figure 3d, 2a-b). Since the region affected by the L340R mutation is restrictive, we concluded that the disruption of interaction was caused by the structural change in the dimer interface. While the key factor would be either or both the C-terminal helix itself and the surface newly formed via dimerization, it is clear that the dimer interface of FIH-1 plays a critical role, namely in terms of contribution to the Δ*G* in the interaction with Mint3.

### Regions of FIH-1 involved in the interaction

Oxygen-dependent regulation of HIF-1 activity is known to be mediated by the hydroxylation of Asn803 at its HIF-1α subunit. This modification is accomplished by the catalytic activity of FIH-1 as an asparaginyl hydroxylase. FIH-1 has a catalytic center, Fe(II), in its β-strand jellyroll core with its co-substrate αKG. Hence, FIH-1 is categorized into the Fe(II)- and αKG-dependent dioxygenases (15). Since previous research suggested that Mint3 inhibits the regulatory activity of FIH-1 by competing with HIF-1α in the PPI, we hypothesized that the catalytic pocket, an obvious binding site of HIF-1α, may be involved in the interaction with Mint3 (3). First, ITC measurements were performed to examine the effect of the co-factors on the interaction between FIH-1 and Mint3NT. The absence of Fe(II) and αKG reduced the Δ*G* by 0.5 and 0.7 kcal.mol^-1^, respectively, and the decrease in Δ*G* caused by the elimination of both co-factors (1.3 kcal.mol^-1^) was approximately the sum of each value (Figure 4a-b). According to the values, each co-factor is not critical but is certainly involved in the interaction. In addition, as the effect on Δ*G* is additive, it is likely that each co-factor independently affects the binding affinity. Moreover, αKG obviously influenced the Δ*H*, whereas Fe (II) did not (Figure 4a-b). Therefore, we assumed that the co-factors, especially αKG, may stabilize the conformation of FIH-1 that binds to Mint3.

**Figure 4:**
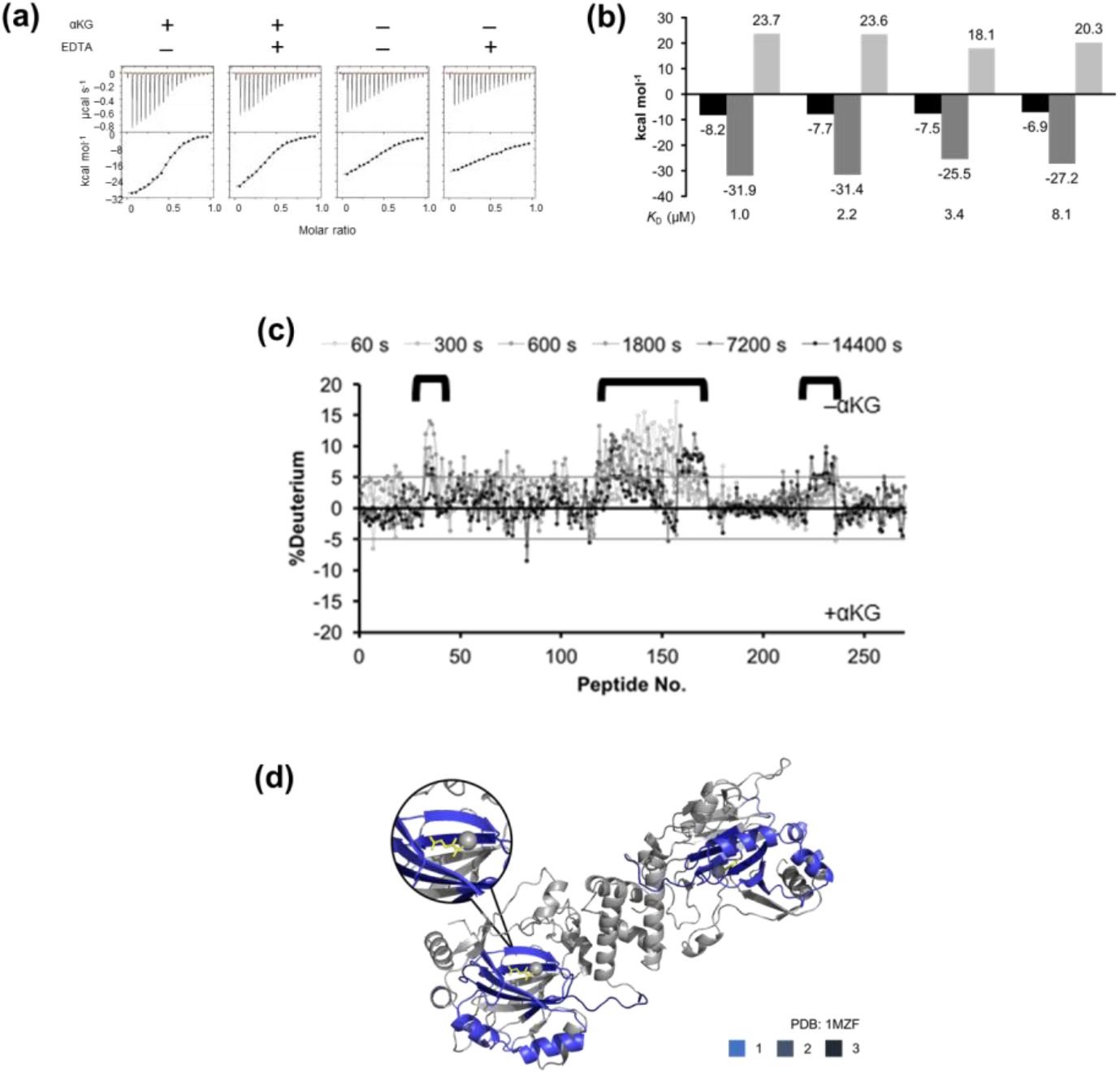
**(A)** ITC profiles and **(B)** thermodynamic parameters of the interaction between FIH-1 and Mint3NT, in the presence/absence of αKG and/or EDTA. **(C)** Differences between HDX-MS charts for Mint3NT, in the absence and presence of αKG. **(D)** Structural image of FIH-1, with the differences in HDX-MS data shown in blue (PDB ID: 1MZF).

Next, HDX-MS was performed against FIH-1 ± αKG, to further investigate its effect on the structure of FIH-1. Upon comparing the deuteration levels of the apo- and holo-forms of FIH-1, a significant difference was observed in several regions (Figure 4c). Merging the results of HDX-MS and the crystal structure revealed that some of the varied regions were around the binding site of αKG, namely the pre-described catalytic pocket. However, it was also discovered that αKG affects not only its binding site, but also its adjacent helical region located at the outer edge of the FIH-1 dimer (Figure 4d). Subsequently, HDX-MS was conducted against FIH-1 ± Mint3NT, to detect the binding site. As a result, the region in FIH-1, the deuteration level of which was shown to be affected by the presence of Mint3NT, corresponded to the αKG-related site (Figure 5a-b). Accordingly, the corresponding region, namely the outer helices (hereafter called the HDX region), is likely to be involved in the interaction with Mint3NT. Meanwhile, it should be noted that there was no information about the dimer interface because it is buried and intrinsically hard to deuterate.

**Figure 5:**
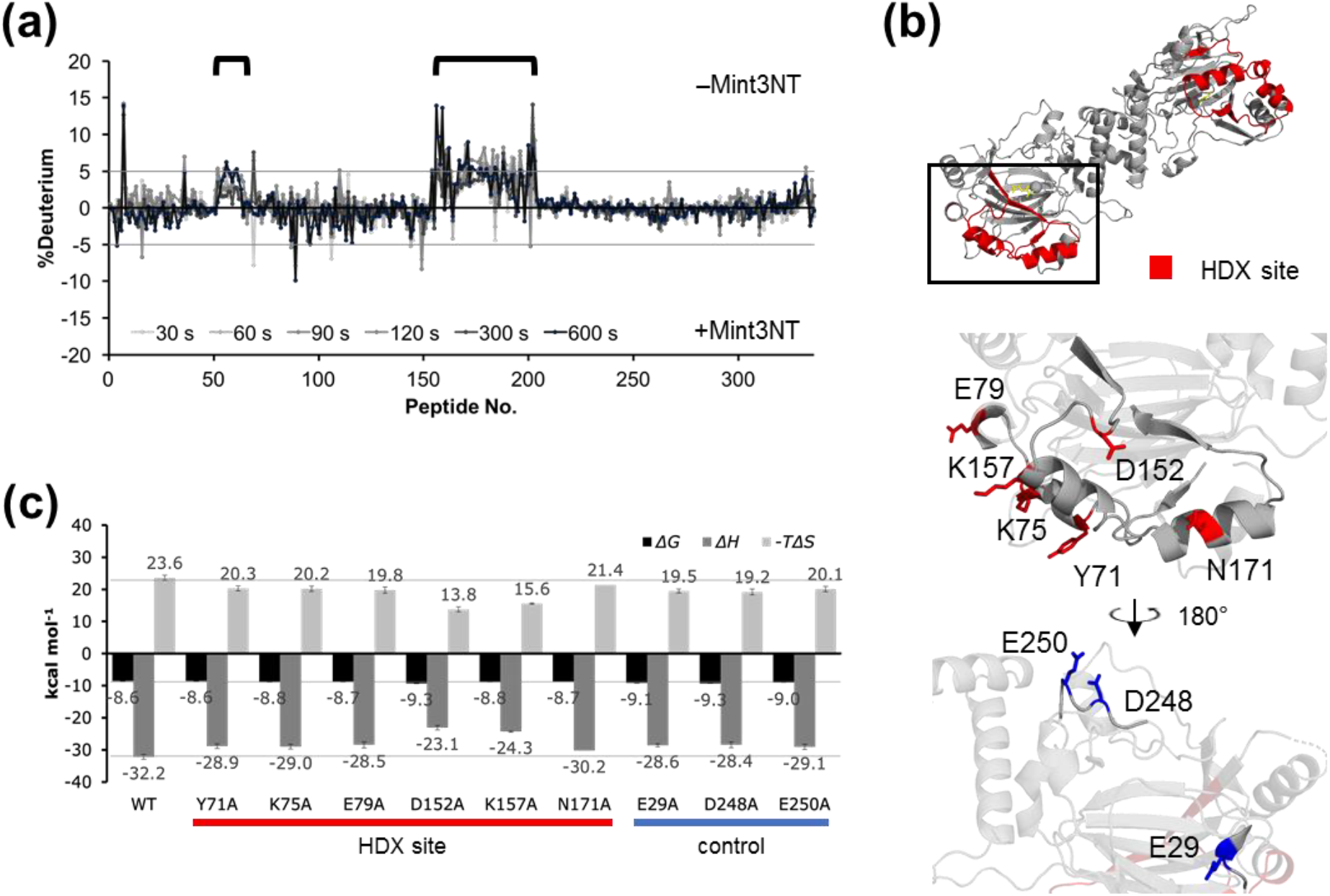
**(A)** Differences between HDX-MS charts for FIH-1 in the absence and presence of Mint3NT. **(B)** Mutation sites of FIH-1 (top: whole structure showing HDX areas in red; middle: structural image focusing on the red area; bottom: structural image focusing on the blue area; PDB ID: 1MZF). **(C)** Thermodynamic parameters of the interactions between FIH-1 and mutations of Mint3NT.

To validate the role of the HDX region, mutation analysis was performed using ITC. Six amino acids within the HDX region (Tyr71, Lys75, Glu79, Asp152, Lys157, and Asn171) were selected based on their orientation and polarity, and replaced with Ala. In addition, three amino acids (Glu29, Asp248, and Glu250) apart from the HDX region were replaced with Ala, as negative controls (Figure 5b). The thermodynamic parameters of binding to Mint3NT were measured three times for each FIH-1 mutant, and the standard deviation was calculated as the experimental error. In terms of the *ΔG*, there was no significant difference between the mutants, including the negative controls. On the other hand, the Δ*H* commonly decreased by approximately 3 kcal mol^-1^ for most of the mutants. According to the result of the negative control, the ∼3 kcal mol^-1^ decrease in Δ*H* was probably caused by the mutation itself. However, D152A and K157A showed a significant decrease of approximately 8 kcal mol^-1^, which was obviously larger than 3 kcal mol^-1^ (Figure 5b-c). Notably, the CD spectra of each mutant suggested that even a single mutation in FIH-1 may affect its structure, but the magnitude of the structural change did not correlate with the decrease in Δ*H* (Figure S7). Therefore, a common decrease in Δ*H* can be expressed by the structural fluctuations. However, the Δ*H* decrease caused by D152A and K157A should be interpreted in terms of its interaction with Mint3NT. In conclusion, the HDX region of FIH-1 is involved in the interaction with Mint3 by contributing not to Δ*G*, but to Δ*H*.

### Identification of the other binding site in Mint3NT

Although the 101-110 region was confirmed to be a pivotal binding site, there is presumably another binding site that exists within the N-terminal half (1-117) of Mint3NT. To identify the other predicted binding sites, we performed HDX-MS for isolated Mint3NT (apo-form) and Mint3NT-FIH-1 complex (holo-form), since the HDX-MS experiment can provide structural information in peptide-scaled resolution. First, to examine whether the HDX-MS system is feasible for detecting the binding site of Mint3NT, the results of the peptides that contain the known binding site (101-110) were used as a positive control. For instance, as expected, the peptides IADAH<u>GLLSAEAGRD</u>DL (96-112) and <u>AGRD</u>DLLGL (107-115) exhibited a significant difference in the deuteration level between the apo- and holo-states. Moreover, the apo- and holo-states of the peptide PEEPPAGAQSPE (184-195), which is located in the C-terminal half of Mint3NT, did not differ in terms of deuteration levels. This result is consistent with the speculation that the C-terminal half of Mint3NT does not contribute to the binding to FIH-1 (Figure 6a). To confirm system reliability, other peptides within the N-terminal half of Mint3NT were also investigated. An obvious difference in the deuteration level between the apo- and holo-states was observed for the peptides lying immediately before the known binding site, for example, LVGPSPGGAPCPLHIATGHGLASQE (71-95) (Figure 6a). To determine the exact region that is important for binding, the amino acid sequence of human Mint3 was compared with that of mouse and rat. Consequently, a well conserved region (78-88) was found within the peptide of concern. Following that, a new mutant of Mint3NT was constructed by altering five amino acids with a large side chain to alanine within the species-common region (hereafter called Mint3mut2) (Figure 6b). In the binding analysis using ITC, a remarkable decrease in binding affinity was observed for Mint3mut2, as compared to Mint3NT (Figure 6c). In addition, a short fragment (78-117) of Mint3NT containing both the detected binding sites was constructed. Compared to the pre-described peptide 93-117, there was an increase in the binding affinity of peptide 78-117, owing to the presence of a newly detected binding site (Figure 6d-e). The attenuation of the binding to FIH-1 caused by the mutation introduced to 78-88 in full-length Mint3 (Mint3mut2) was also observed in a cellular assay using immunoprecipitation (Figure 7a). In addition, the activation effect of HIF-1α mediated via the interaction between Mint3 and FIH-1 was significantly disrupted by this mutation (Figure 7b). Accordingly, we concluded that the 78-88 region is a major binding site in Mint3NT.

**Figure 6:**
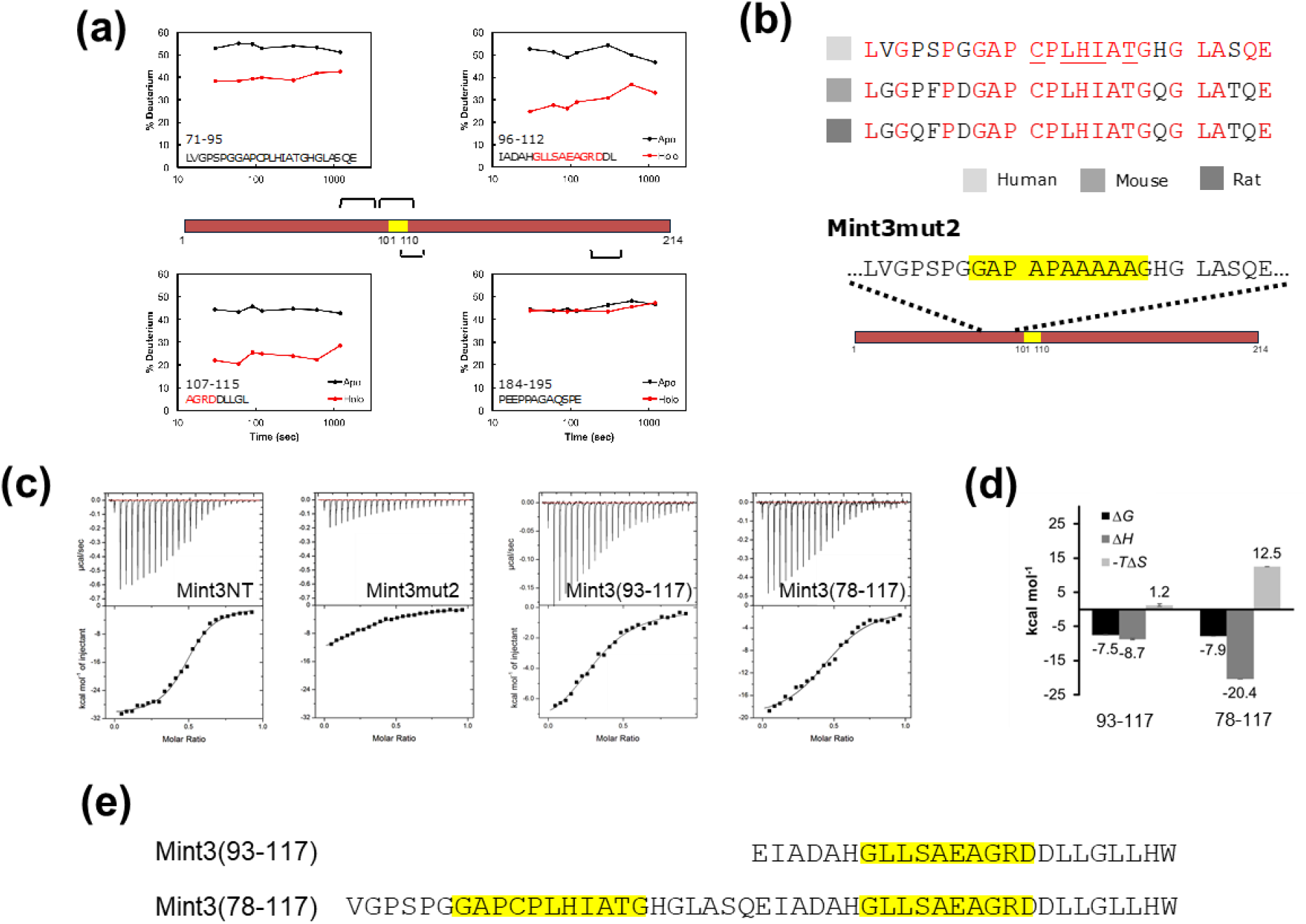
**(A)** HDX-MS profiles of Mint3NT peptides, in the absence and presence of FIH-1. **(B)** Comparison of the amino acid sequence of Mint3NT among animal species, and amino acid sequence of Mint3mut2. **(C)** ITC profiles of the interaction between FIH-1 and Mint3NT mutants. **(D)** Thermodynamic parameters of Mint3(93-117) and Mint3(78-117). **(E)** Amino acid sequences of Mint3(93-117) and Mint3(78-117).

**Figure 7:**
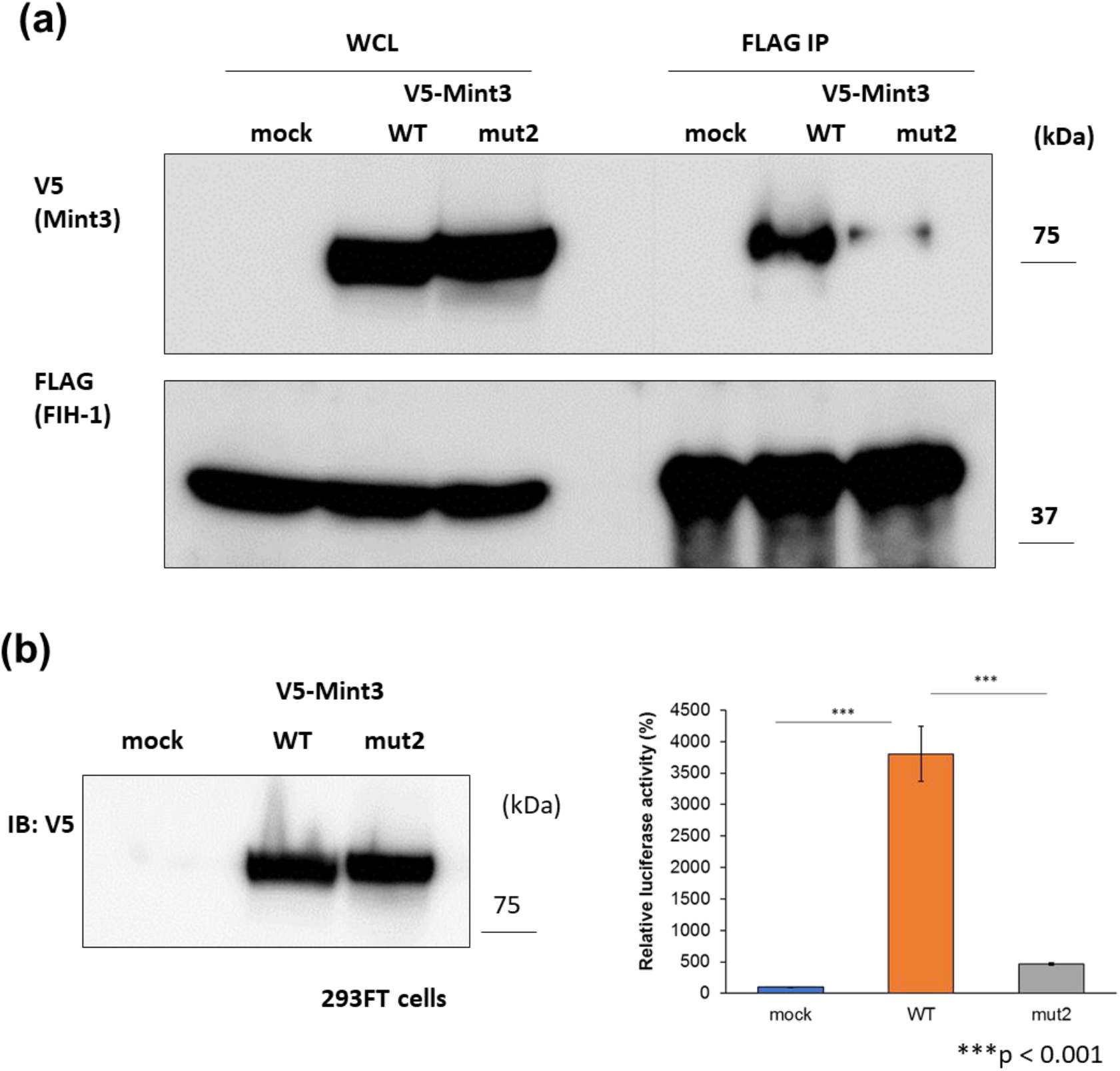
**(A)** Immunoblotting using mouse anti-V5 antibody and rabbit anti-FLAG polyclonal antibody in 293FT cells. Left side is a sample of the whole cell lysate, while right side is the sample immunoprecipitated using an anti-FLAG polyclonal antibody. **(B)** Reporter assay expressed either vector alone, wild-type Mint3, or Mint3mut2 in 293FT cells. Left side is the immunoblotting data of the whole cell lysate using an anti-V5 antibody, while right side is the data for the luciferase activities.

### Description of the Mint3-FIH-1 complex

Through thermodynamic analysis using ITC and structural analysis using HDX-MS, we made four main observations: (i) Mint3 has two core regions (77-87 and 101-110), which contribute to its *ΔG* to FIH-1. (ii) The interfacial structure of the FIH-1 dimer plays a critical role in the interaction with Mint3. (iii) The regions of Mint3 other than the cores are not involved in the Δ*G*, while they do affect the Δ*H*. (iv) Some amino acids (Asp152 and Lys157) in the outer helices of the FIH-1 dimer do not contribute to the Δ*G*, but contribute to the Δ*H*. Based on these results, we generated a model of the Mint3-FIH-1 complex, composed of “binding” part and “touching” parts (Figure 8a). Here, the “binding” parts indicate the parts contributing to the Δ*G*, and “touching” parts represent the parts involved in the interaction by affecting the *ΔH*, rather than the Δ*G*. In the “binding” part, the core regions (77-87 and 101-110) of Mint3 are interacting with the dimer interface of FIH-1, reflecting their critical roles in the Δ*G*. On the other hand, the “touching” parts are composed of 1-76/111-214 of Mint3 and the outer helices of FIH-1 dimer, indicating that both of them contribute to produce balanced favorable enthalpy and unfavorable entropy through the interaction.

**Figure 8:**
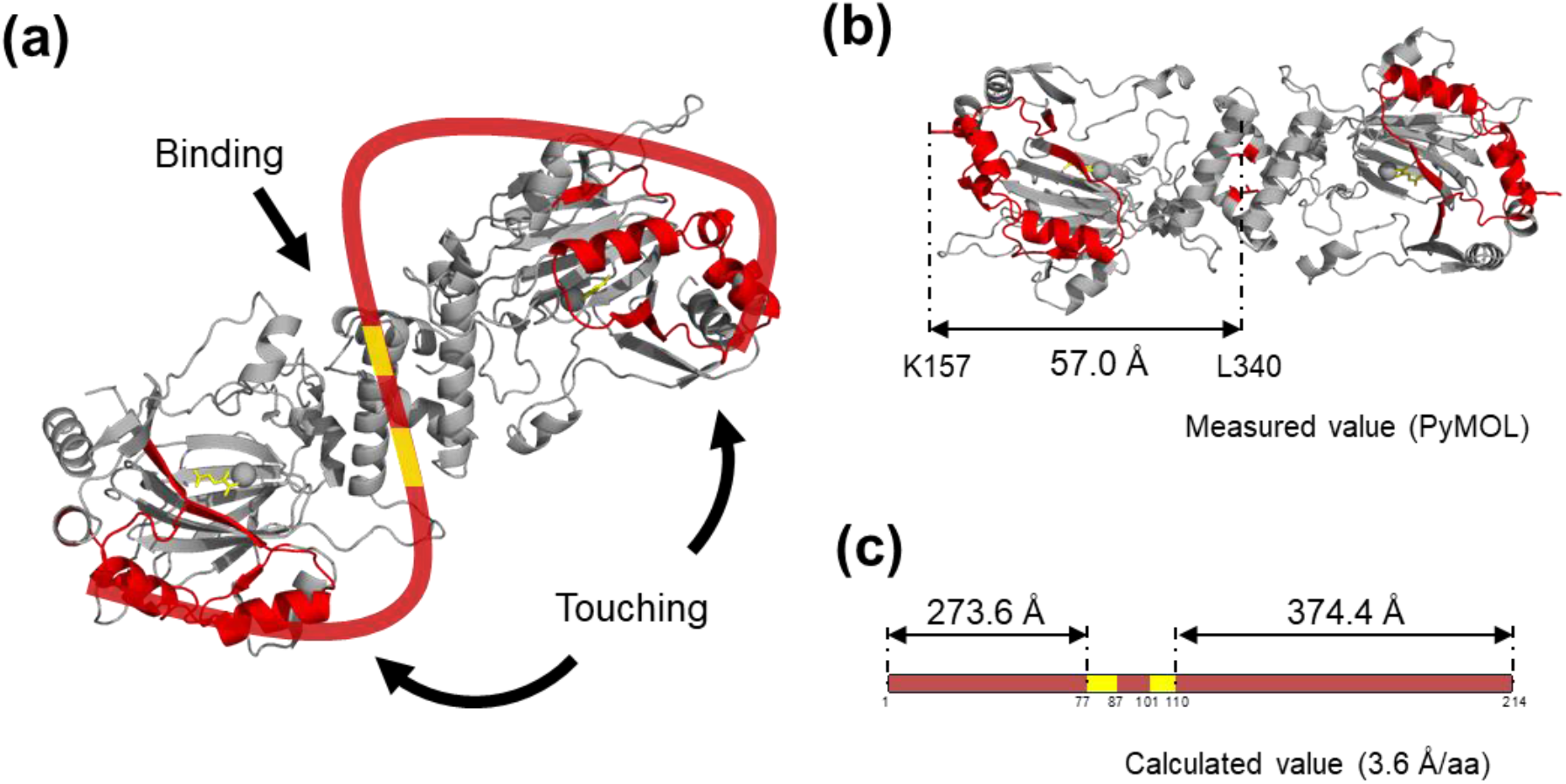
**(A)** The interaction model between FIH-1 and Mint3NT, as inferred from this study. The three-dimensional structure is that of FIH-1 dimer, while the string-like illustration is a schematic for Mint3NT (PDB ID: 1MZF). **(B)** The size of the FIH-1 dimer was calculated using Pymol software (PDB ID: 1MZF). **(C)** The estimated length of Mint3NT when extended on a linear chain.

To confirm that the description was physically possible, we estimated the molecular size of the proteins. The scale of FIH-1 was obtained from its crystal structure, by measuring the distance between the outer helix (Lys157) and the center of the dimer interface (Leu340), which represents the semi-major axis of the FIH-1 dimer. As a result, the value was estimated at 57.0 Å (Figure 8b). Meanwhile, the linear length of Mint3NT was calculated from the average end-to-end distance of one amino acid (3.6 Å) (16–19). Consequently, the lengths of 1-76 and 111-214 regions of Mint3NT were estimated at 273.6 Å and 374.4 Å, respectively (Figure 8c). This model is consistent with the absence of secondary structural changes in the CD analysis. According to the results, Mint3 is long enough to “bind” to the dimer interface and “touch” at the outer helix of the FIH-1 dimer simultaneously.

## Discussion

Mint3 is a member of the X11 protein family that has conserved C-terminal domains, such as the binding site of APP and unique N-terminal domains (20, 21). Previous research has demonstrated that the N-terminal half of Mint3 binds to FIH-1 and acts as an activator of the HIF-1 pathway, which is involved in cellular responses to hypoxia and leads to the proliferation or metastasis of cancer cells (4, 13). However, the conformation and mechanism of the interaction between Mint3 and FIH-1 remain unclear. Thus, using CD, HDX-MS, and NMR analyses, we first demonstrated that the N-terminal portion, which is the functional part of Mint3, is highly disordered in its isolated form. This result is consistent with the emerging knowledge of IDRs. In particular, their biological roles include signal transfer, regulation, transcription, and replication (22, 23).

In general, many IDRs undergo transitions to more ordered or folded states upon binding to their targets, to execute their biological functions. The kinase-inducible transcriptional activation domain (KID) of cyclic AMP response element-binding protein (CREB) is one of the most characterized examples (22, 24, 25). Although KID is intrinsically disordered, it forms an orthogonal helix by binding to the CREB-binding protein (CBP) (26). However, although the unfavorable entropy suggested a loss of flexibility in binding to FIH-1, coupled folding and binding could not be observed upon CD measurement of Mint3NT. Therefore, we hypothesized that there is a distinct mechanism for PPI between Mint3NT and FIH-1.

To reveal the binding mechanism, we conducted thermodynamic and structural analyses using ITC and HDX-MS, respectively, since the classical approaches, particularly crystallographic methods, have little relevance for a mostly disordered protein such as Mint3NT. Although these methods can not directly visualize the interaction, we succeeded in identifying the regions involved in the interaction, “binding” and “touching” sites, and revealing their different roles in the thermodynamics, as mentioned above (Figure 8a). The “binding” sites of the dimer interface of FIH-1 for 78-88 and 101-110 regions of Mint3NT, which represent the regions that contribute to the Δ*G*, is a usual interpretation. In particular, we proposed that the 78-88 amino acids in Mint3 are a critical binding region to FIH-1. On the other hand, the “touching” sites, which indicate the parts that do not contribute to the Δ*G* but certainly influence the Δ*H*, is somewhat conceptual. Some previous reports can provide hints for interpreting the roles of the “touching” site. Tummino et al. suggest enhancing the enthalpic contributions of protein interactions would likely favor slower dissociation and, hence, a longer residence time (27). In addition, it has been reported IDPs/IDRs can regulate the binding kinetics (28). Therefore, the “touching” sites may affect the rate constant of the interaction, although further investigation is required in this regard.

Our model for the complex of Mint3NT and FIH-1 mentioned above is a description that can consistently explain all the results. Notably, the region between the “binding” site and the “touching” site of Mint3 is delineated to cover the biding pocket of αKG in FIH-1. This model is consistent with the fact that Mint3 abrogates the ability of FIH-1 to modify HIF-1α, by competing with HIF-1α to bind to the αKG binding site of FIH-1 (3). Although the detailed conformation of Mint3 needs to be determined directly using some experimental methods, this model can provide us insight into the suppression mechanism of Mint3 toward HIF-1 binding to FIH-1.

In addition, the detailed binding mechanism can be used as a guide for drug development in cancer therapy. According to the Mint3-FIH-1 complex model, there are two possible strategies for disturbing the suppression activity of Mint3. The first strategy is to aim at a critical site of binding affinity, which in this case is the “binding” site, as it seems to be more effective in inhibiting the interaction with FIH-1. The second strategy is to focus on the “touching” site, to prevent Mint3 from covering the αKG-binding site. The merit of the second scenario is that the inhibitor with lower affinity may still be effective in this case, as compared to the case with aiming at the “binding” site.

In conclusion, we found that the N-terminal region of Mint3 has properties corresponding to IDRs, and that this region interacts with FIH-1 in an enthalpy-driven manner. Using Mint3nt mutants, we suggest that the core region of Mint3 binds to FIH-1 in such a way that a single molecule rests on the FIH-1 dimer interface. We believe that the binding mechanism underlying FIH-1 suppression caused by Mint3 has been elucidated by our thermodynamic and structural analyses, and this insight can serve as a foothold for further investigation of IDRs and drug development for cancer therapy.

## Experimental Procedures

### Recombinant protein expression and purification

The coding sequence for human Mint3NT (1-214) with a C-terminal hexa-histidine (His_6_) tag, and human FIH-1 (full-length) with an N-terminal His_6_ tag and cleavage site (ENLYFQ) of TEV protease were inserted into the pET28 vector. The plasmids were transformed into BL21 (DE3) cells and cultured at 28°C in Luria broth medium containing 50 μg/mL kanamycin. M9 minimal medium containing 0.5 g/L ^15^NH_4_Cl was used for the preparation of uniformly ^15^N-labeled protein. Protein synthesis was induced by the addition of 1 mM Isopropyl β-D-1-thiogalactopyranoside (IPTG) to the culture, when its optical density at 600 nm reached a value of ∼0.6, and further culturing at 20°C for 16 h. Cells were harvested by carrying out a round of centrifugation at 7000 × *g* for 10 min and disrupted by sonication in a binding buffer composed of 20 mM Tris-HCl (pH 8.0), 500 mM NaCl, and 5 mM imidazole. The lysates were centrifuged at 40000 × *g* for 30 min, following which the supernatants were loaded onto a Ni-NTA Agarose (Qiagen, Hilden, Germany) column that had been pre-equilibrated with the binding buffer. The column was washed with 20 column volumes of wash buffer composed of 20 mM Tris-HCl (pH 8.0), 500 mM NaCl, and 20 mM imidazole. His _6_-tagged Mint3NT and His_6_-tagged FIH-1 were eluted with 10 mL of an elution buffer composed of 20 mM Tris -HCl (pH 8.0), 500 mM NaCl, and 200 mM imidazole. The His_6_ tag of FIH-1 was cleaved using His_6_-tagged TEV protease. Mint3NT was purified using SEC with a HiLoad 26/60 Superdex^®^ 200 pg or 75 pg column (GE Healthcare, Illinois, United States) at 4°C. The monomer fraction was collected from the purification. FIH-1 was purified using SEC with HiLoad 26/60 Superdex^®^ 200 pg at 4°C. The elution profiles of both the proteins were monitored at a wavelength of 280 nm. The mutants/fragments of Mint3 and FIH -1 were prepared as described above.

### Circular dichroism analysis

Far ultraviolet (UV) CD spectra of Mint3NT, FIH-1, and FIH-1 mutants were measured on a J-820 spectropolarimeter (Jasco, Tokyo, Japan) at 25°C in 35 mM Na-phosphate (pH 7.5), 300 mM NaCl, and 500 μΜ αKG. The concentrations of the protein samples were maintained in the range of 5-10 μΜ, as appropriate to keep the photomal voltage under 600 V. Each spectrum was generated upon accumulation of 5 measurements.

### Hydrogen/deuterium exchange mass spectrometry analysis

Mint3NT, FIH-1, and its mutants were prepared in Na-phosphate buffer composed of 35 mM Na-phosphate (pH 7.5) and 300 mM NaCl at a final concentration of 1.0 mg mL^-1^. Each protein was diluted 10-fold with Na-phosphate buffer (the solvents are as mentioned before) in Heavy water (D_2_O). The diluted solutions were then incubated separately at 10°C. Deuterium-labeled samples were quenched by diluting about 10-fold with quenching buffer at pH 2.3, composed of 2 M guanidine hydrochloride and 200 mM citric acid. All of the processes described above were performed automatically using HDX Workflow Solution (LEAP Technologies, Bjerndrup, Denmark), and all the time-points were determined in three independent labeling experiments. After quenching, the solutions were subjected to online pepsin digestion followed by LC/MS analysis using an UltiMate™ 3000 RSLCnano System (Thermo Fisher Scientific, Massachusetts, United States) connected to a Q Exactive™ Plus mass spectrometer (Thermo Fisher Scientific). Online pepsin digestion was performed with a Poroszyme ^®^ Immobilized Pepsin Cartridge 2.1 × 30 mm (Waters Corporation, Massachusetts, United States) in formic acid solution (pH 2.5) for 3 min at 8°C, at a flow rate of 50 µL min^-1^. Desalting and analytical processes after pepsin digestion were performed using Acclaim ™ PepMap™ 300 C18 (1.0 × 15 mm; Thermo Fisher Scientific) and Hypersil GOLD™ (1.0 × 50 mm; Thermo Fisher Scientific) columns. The mobile phase was 0.1% formic acid solution (buffer A) and 0.1% formic acid containing 90% acetonitrile (B buffer). The deuterated peptides were eluted at a flow rate of 45 μL min^-1^, with a gradient of 10%–90% of buffer B for 9 min. The conditions of the mass spectrometer were as follows: electrospray voltage, 3.8 kV; positive ion mode, sheath and auxiliary nitrogen flow rate at 20 and 2 arbitrary units; ion transfer tube temperature at 275°C; auxiliary gas heater temperature at 100°C; and a mass range of m/z 200–2,000. Data-dependent acquisition was performed using a normalized collision energy of 27 arbitrary units. Analysis of the deuteration levels of the peptide fragments was performed based on the MS raw files, by comparing the spectra of deuterated samples with those of non -deuterated samples, using the HDExaminer software (Sierra Analytics, California, United States).

### Nuclear magnetic resonance spectroscopy

^1^H-^15^N HSQC spectra were measured on an Avance™ I 600 spectrometer (Bruker BioSpin, Switzerland) equipped with a TXI-triple resonance triple gradient probe at 4°C. The sample concentration was 50 μM ^15^N-labeled protein dissolved in 50 mM Na-phosphate buffer (pH 7.3) containing 10% D_2_O. The spectra were processed using NMRPipe (29).

### Differential scanning calorimetry analysis

Thermal denaturation experiments were performed using a MicroCal™ VP-DSC calorimeter (GE Healthcare). Mint3NT was dialyzed against Na-phosphate buffer composed of 35 mM Na-phosphate (pH 7.5), 300 mM NaCl at 4°C for 16 h, and concentrated to 50 μΜ (∼1.2 mg mL^- 1^) using Amicon^®^ Ultra-15 Centrifugal Filter Units (EMD Millipore). The protein was loaded into the sample cell and heated from 10°C to 100°C, at 60°C h^-1^. The thermogram was generated using Origin 7 (OriginLab, Massachusetts, United States).

### Isothermal titration calorimetry

Calorimetric titration of Mint3NT with FIH-1 was performed using a MicroCal™ iTC_200_ system (GE Healthcare) at 25°C. Both proteins were dialyzed against Na-phosphate buffer composed of 35 mM Na-phosphate (pH 7.5), 300 mM NaCl, and 500 μΜ αKG at 4°C for 16 h and concentrated using Amicon^®^ Ultra-15 Centrifugal Filter Units. At each injection (total 25 times), 1.5 μL of 150 μM Mint3NT solution was added to a sample cell containing 30 μM FIH-1 solution. All titration data were analyzed by fitting to a single set of site models using Origin 7 (OriginLab, Massachusetts, United States). Calorimetric titration of Mint3NT mutants/fragments and FIH-1 mutants was performed as described above.

### SEC-multi-angle laser scattering

MALS was detected using DAWN8+^®^ (Wyatt Technology, California, United States). FIH-1 was dialyzed against Na-phosphate buffer composed of 35 mM Na-phosphate (pH 7.5), 300 mM NaCl, and 500 μΜ αKG at 4°C for 16 h, and concentrated to 70 μM using Amicon^®^ Ultra-15 Centrifugal Filter Units. The protein sample (50 μL) was loaded into a 10/300 Superdex^®^ 200 GL (GE Healthcare) column connected to the MALS detection unit for SEC. During the experiment, UV detection was performed using an SPD-10A UV/Vis Detector (Shimadzu, Kyoto, Japan). The obtained data were processed and analyzed using ASTRA (Wyatt Technology).

### Differential scanning fluorometry analysis

The thermal shift assay was performed on a CFX Connect™ Real-Time System (Bio-Rad, California, United States). FIH-1 wild-type and L340R were dialyzed against Na-phosphate buffer composed of 35 mM Na-phosphate (pH 7.5), 300 mM NaCl, and 500 μΜ αKG at 4°C for 16 h, and concentrated to 10 μM using Amicon^®^ Ultra-15 Centrifugal Filter Units. One thousandth of the protein sample volume of SYPRO ™ Orange Protein Gel Stain (5000 × concentrated in Dimethyl sulfoxide (DMSO); Invitrogen, California, United States) was added, and the proteins containing 5× SYPRO™ Orange were loaded onto Hard-Shell^®^ 96-well PCR Plates (Bio-Rad) for measurement. The denaturation of each protein was observed via fluorescence emitted from SYPRO ™ Orange upon absorption to the exposed hydrophobic surface.

### Cellular analysis using immunoprecipitation and reporter assay

293FT cells, which are derived from HEK293 human embryonic kidney cells and express the simian virus large T antigen, were purchased from Thermo Fisher Scientific. Cells were cultured at 37°C in a humidified atmosphere containing 5% CO_2_ in high-glucose Dulbecco’s Modified Eagle Medium (DMEM) (Thermo Fisher Scientific) containing 10% fetal bovine serum, 100 units/mL penicillin, and 100 µg/mL streptomycin (Merck, Massachusetts, United States).

pcDNA3 (Thermo Fischer Scientific) expression vectors for V5-tagged Mint3 and FLAG-tagged FIH-1 were constructed as previously described (3). Expression vectors expressing mutant Mint3 (Mint3mut2) were generated using PCR-based methods.

Immunoprecipitation was performed as previously described (3, 30). Briefly, 293FT cells were co-transfected with expression plasmids encoding a V5-tagged Mint3 construct and a FLAG-tagged FIH-1 construct using Lipofectamine™ 2000 (Thermo Fischer Scientific). Cells were lysed in lysis buffer 24 h after transfection and centrifuged at 20,000 × *g* for 15 min at 4°C. Supernatants were collected and incubated with beads conjugated to anti-FLAG M2 antibody (Merck). Beads were washed, and the proteins bound to them were eluted using FLAG peptide and analyzed using immunoblotting. Immunoblotting was performed as previously described (3, 30) using mouse anti-V5 antibody (R960-25, Thermo Fisher Scientific) and rabbit anti-FLAG polyclonal antibody (F7425, Merck).

Reporter assays were performed as previously described, with minor modifications (3, 31). A pGL4.35 reporter vector containing the firefly luciferase gene, under the control of a transcriptional regulatory unit comprising 9× Gal4-binding elements, was purchased from Promega, Wisconsin, United States. A pRL vector expressing Renilla luciferase (Promega) served as an internal control. 293FT cells were seeded into 24-well plates at a density of 2.5×10^4^ cells/well and co-transfected with a reporter plasmid (100 ng), internal control vector (10 ng), pcDNA3 Gal4BD-HIF-1α CAD plasmid (3, 31) (50 ng), and other plasmids (200 ng) that expressed either vector alone, wild-type Mint3, or Mint3mut2. Transfection was performed using Lipofectamine™ 2000. Twenty-four hours after transfection, luciferase activity was measured using the Dual-Glo^®^ Luciferase Assay System (Promega), according to the manufacturer’s instructions. Luminescence was measured using a GloMax^®^ 20/20 luminometer (Promega).

## Acknowledgments and Funding Information

We thank Thermo Fisher Scientific for the technical support in HDX-MS experiments. This work was supported by JSPS KAENHI under grant number JP18H02082, JP18H05425 (to S.N.) and JP16H02420, JP19H05766, JP20H02531 (to K.T.), by the Platform Project for Supporting Drug Discovery and Life Science Research [Basis for Supporting Innovative Drug Discovery and Life Science Research (BINDS)] from AMED of Japan under grant number JP20am0101094 (to K.T.), and by the P-CREATE (Project for Cancer Research and Therapeutic Evolution) from AMED of Japan under grant number 16770655 (to S.N., T.S., and K.T.).

## Footnotes

This article contains supporting information.

## Abbreviations

FIH-1: factor inhibiting HIF -1;
αKG: α-ketoglutaric acid;
CD: circular dichroism;
DSC: differential scanning calorimetry;
HDX-MS: hydrogen/deuterium exchange mass spectrometry;
NMR: nuclear magnetic resonance;
ITC: isothermal titration calorimetry;
SEC: size-exclusion chromatography;
SEC-MALS: SEC-multi-angle laser scattering;
DSF: differential scanning fluorometry

## References

1. DeBerardinis, R. J., and Chandel, N. S. (2020) We need to talk about the Warburg effect. Nat Metab. 2, 127–129

2. Potter, M., Newport, E., and Morten, K. J. (2016) The Warburg effect: 80 years on. Biochem. Soc. Trans. 44, 1499–1505

3. Sakamoto, T., and Seiki, M. (2009) Mint3 enhances the activity of hypoxia-nducible factor-1 (HIF-1) in macrophages by suppressing the activity of factor inhibiting HIF -1. J. Biol. Chem. 284, 30350–30359

4. Uversky, V. N. (2014) Introduction to intrinsically disordered proteins (IDPs). Chem. Rev. 114, 6557–6560

5. Liu, Y., Wang, X., and Liu, B. (2019) A comprehensive review and comparison of existing computational methods for intrinsically disordered protein and region prediction. Brief. Bioinform. 20, 330–346

6. Csizmok, V., Follis, A. V., Kriwacki, R. W., and Forman-Kay, J. D. (2016) Dynamic protein interaction networks and new structural paradigms in signaling. Chem. Rev. 116, 6424–6462

7. Tompa, P., Schad, E., Tantos, A., and Kalmar, L. (2015) Intrinsically disordered proteins: emerging interaction specialists. Curr. Opin. Struct. Biol. 35, 49–59

8. Okamoto, M., and Südhof, T. C. (1998) Mint 3: a ubiquitous mint isoform that does not bind to munc18-1 or -2. Eur. J. Cell Biol. 77, 161–165

9. Hata, Y., Slaughter, C. A., and Südhof, T. C. (1993) Synaptic vesicle fusion complex contains unc-18 homologue bound to syntaxin. Nature. 366, 347–351

10. Kelly, S. M., Jess, T. J., and Price, N. C. (2005) How to study proteins by circular dichroism. Biochim. Biophys. Acta. 1751, 119–139

11. Konermann, L., Pan, J., and Liu, Y. -H. (2011) ChemInform abstract: Hydrogen exchange mass spectrometry for studying protein structure and dynamics. ChemInform. 42, no-no

12. Hamuro, Y., Coales, S. J., Morrow, J. A., Molnar, K. S., Tuske, S. J., Southern, M. R., and Griffin, P. R. (2006) Hydrogen/deuterium -exchange (H/D-Ex) of PPARgamma LBD in the presence of various modulators. Protein Sci. 15, 1883–1892

13. Sakamoto, T., Niiya, D., and Seiki, M. (2011) Targeting the Warburg effect that arises in tumor cells expressing membrane type-1 matrix metalloproteinase. J. Biol. Chem. 286, 14691–14704

14. Lancaster, D. E., McNeill, L. A., McDonough, M. A., Aplin, R. T., Hewitson, K. S., Pugh, C. W., Ratcliffe, P. J., and Schofield, C. J. (2004) Disruption of dimerization and substrate phosphorylation inhibit factor inhibiting hypoxia-inducible factor (FIH) activity. Biochem. J. 383, 429–437

15. Dann, C. E., 3rd, Bruick, R. K., and Deisenhofer, J. (2002) Structure of factor-inhibiting hypoxia-inducible factor 1: An asparaginyl hydroxylase involved in the hypoxic response pathway. Proc. Natl. Acad. Sci. U. S. A. 99, 15351–15356

16. Ainavarapu, S. R. K., Brujic, J., Huang, H. H., Wiita, A. P., Lu, H., Li, L., Walther, K. A., Carrion-Vazquez, M., Li, H., and Fernandez, J. M. (2007) Contour length and refolding rate of a small protein controlled by engineered disulfide bonds. Biophys. J. 92, 225–233

17. Dietz, H., and Rief, M. (2006) Protein structure by mechanical tri-angulation. Proc. Natl. Acad. Sci. U. S. A. 103, 1244–1247

18. Carrion-Vazquez, M., Li, H., Lu, H., Marszalek, P. E., Oberhauser, A. F., and Fernandez, J. M. (2003) The mechanical stability of ubiquitin is linkage dependent. Nat. Struct. Biol. 10, 738–743

19. Oesterhelt, F., Oesterhelt, D., Pfeiffer, M., Engel, A., Gaub, H. E., and Müller, D. J. (2000) Unfolding pathways of individual bacteriorhodopsins. Science. 288, 143–146

20. Miller, C. C. J., McLoughlin, D. M., Lau, K.-F., Tennant, M. E., and Rogelj, B. (2006) The X11 proteins, Abeta production and Alzheimer’s disease. Trends Neurosci. 29, 280– 285

21. Rogelj, B., Mitchell, J. C., Miller, C. C. J., and McLoughlin, D. M. (2006) The X11/Mint family of adaptor proteins. Brain Res. Rev. 52, 305–315

22. Jensen, M. R., Zweckstetter, M., Huang, J.-R., and Blackledge, M. (2014) Exploring free-energy landscapes of intrinsically disordered proteins at atomic resolution using NMR spectroscopy. Chem. Rev. 114, 6632–6660

23. Dyson, H. J., and Wright, P. E. (2005) Intrinsically unstructured proteins and their functions. Nat. Rev. Mol. Cell Biol. 6, 197–208

24. Radhakrishnan, I., Pérez-Alvarado, G. C., Dyson, H. J., and Wright, P. E. (1998) Conformational preferences in the Ser133-phosphorylated and non-phosphorylated forms of the kinase inducible transactivation domain of CREB. FEBS Lett. 430, 317–322

25. Richards, J. P., Bächinger, H. P., Goodman, R. H., and Brennan, R. G. (1996) Analysis of the structural properties of cAMP-responsive element-binding protein (CREB) and phosphorylated CREB. J. Biol. Chem. 271, 13716–13723

26. Radhakrishnan, I., Pérez-Alvarado, G. C., Parker, D., Dyson, H. J., Montminy, M. R., and Wright, P. E. (1997) Solution structure of the KIX domain of CBP bound to the transactivation domain of CREB: a model for activator:coactivator interactions. Cell. 91, 741–752

27. J., T. P., and A, C. R. (2008) Residence Time of Receptor−Ligand Complexes and Its Effect on Biological Function. Biochemistry. 47, 5481–5492

28. J., D., S., G., and P, J. (2014) The binding mechanisms of intrinsically disordered proteins. Phys. Chem. Chem. Phys. 16, 6323–6331

29. Delaglio, F., Grzesiek, S., Vuister, G. W., Zhu, G., Pfeifer, J., and Bax, A. (1995) NMRPipe: a multidimensional spectral processing system based on UNIX pipes. J. Biomol. NMR. 6, 277–293

30. Nakaoka, H. J., Hara, T., Yoshino, S., Kanamori, A., Matsui, Y., Shimamura, T., Sato, H., Murakami, Y., Seiki, M., and Sakamoto, T. (2016) NECAB3 promotes activation of hypoxia-inducible factor -1 during normoxia and enhances tumourigenicity of cancer cells. Sci. Rep. 6, 22784

31. Sakamoto, T., Weng, J. S., Hara, T., Yoshino, S., Kozuka-Hata, H., Oyama, M., and Seiki, M. (2014) Hypoxia-inducible factor 1 regulation through cross talk between mTOR and MT1-MMP. Mol. Cell. Biol. 34, 30–42

